# Invariant errors reveal limitations in motor correction rather than constraints on error sensitivity

**DOI:** 10.1101/189597

**Authors:** Hyosub E. Kim, J. Ryan Morehead, Darius E. Parvin, Reza Moazzezi, Richard B. Ivry

## Abstract

Implicit sensorimotor adaptation is traditionally described as a process of error reduction, whereby a fraction of the error is corrected for with each movement. Here, in our study of healthy human participants, we characterize two constraints on this learning process: the size of adaptive corrections is only related to error size when errors are smaller than 6°, and learning functions converge to a similar level of asymptotic learning over a wide range of error sizes. These findings are problematic for current models of sensorimotor adaptation, and point to a new theoretical perspective in which learning is constrained by the size of the error correction, rather than sensitivity to error.

---

Movement errors are ubiquitous, arising from numerous sources such as motor noise, fatigue, or changes in the environment. A large body of evidence has revealed that the motor system compensates for errors via sensorimotor adaptation^1^. This implicit learning process is thought to be driven by sensory prediction error, the discrepancy between the actual and predicted sensory outcome of a motor command^2–6^. A core issue for models of adaptation has centered on how this error signal is used to modify motor output^7–10^.

In classic models of sensorimotor adaptation, the response to error is assumed to be linear, with trial-by-trial corrections a constant fraction of error size^9,11^. The theoretical foundation for this relationship centers on the delta learning rule, where the weights between putative sensorimotor neurons are updated as a function of the magnitude of the difference between the actual and predicted output^9,12,13^. A standard formulation of this type of model is given by the following state-space equation:

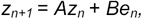

where *z*_*n*_ represents the state estimate of the perturbation on trial *n*, and *A* is a retention factor, the proportion of the state retained from one trial to the next. The error term, *e*, is multiplied by a scalar learning rate, *B*, to determine the change in the state estimate from trial-to-trial due to sensory prediction errors. However, empirical studies have shown limitations with the assumption of a constant learning rate. When operationalized as the ratio of the change in behavior relative to the error, sensitivity appears to be high for small errors, with the system correcting for a relatively large fraction of the error, and then rapidly decreases as errors become large^8,10,14,15^.

However, due to potential confounds in standard sensorimotor adaptation tasks, estimates of the error sensitivity function in many of these studies may be contaminated by other learning processes, such as the use of explicit aiming strategies^16^. To study adaptation without interference from explicit learning or performance-driven corrections, we recently introduced a method in which the visual feedback is task irrelevant and invariant over the course of the experiment^17^ (Fig. 1b). Despite full knowledge of the task-irrelevant clamped visual feedback, participants implicitly produce a marked change in performance. As shown in our initial study with this method, performance changes resulting from clamped feedback bear the classic hallmarks of adaptation, including sign-dependent corrections, persistent aftereffects, local generalization, and a dependency on the integrity of the cerebellum.

**Figure 1.**
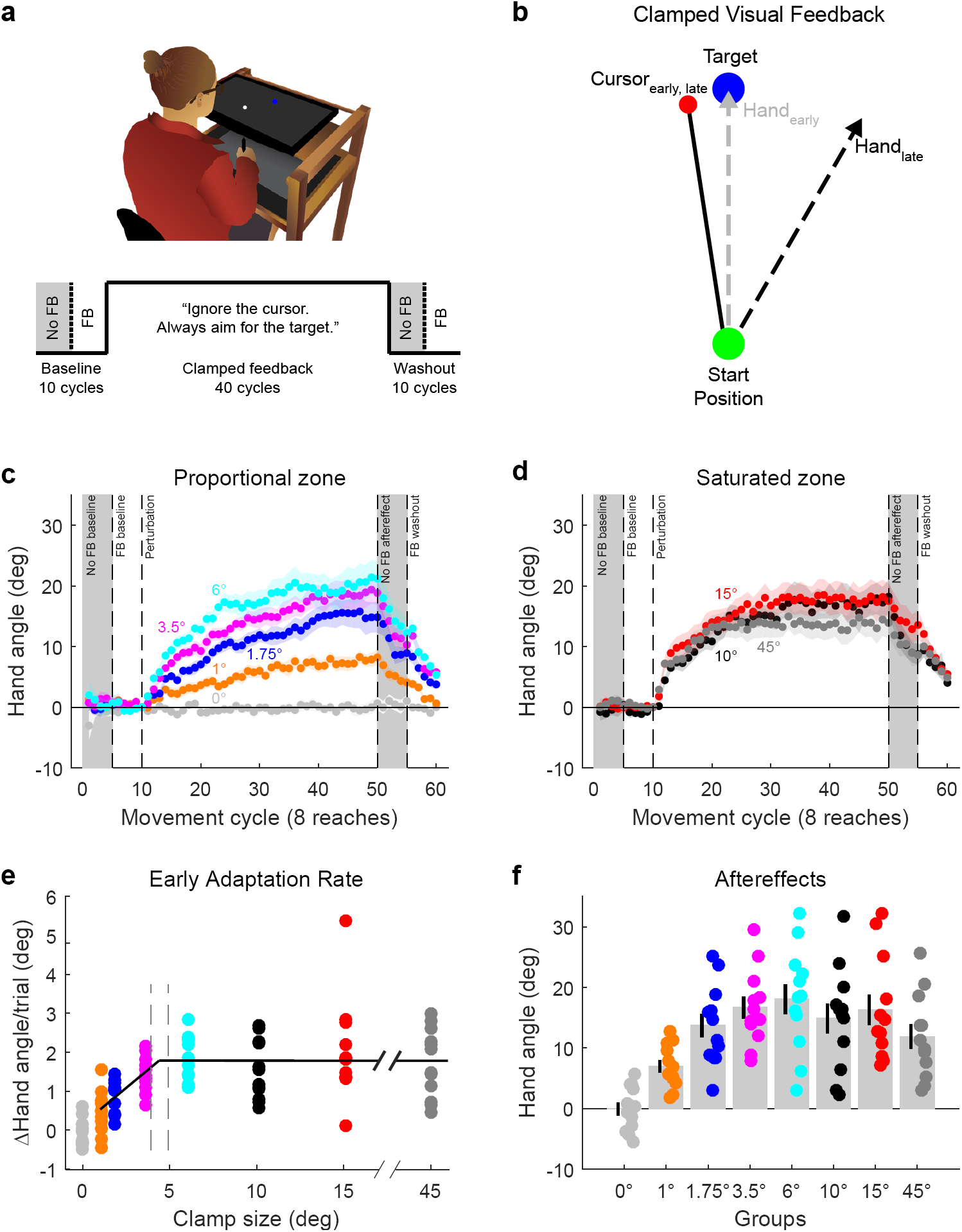
Initial adaptation rates scale with error size, yet saturate to an invariant response magnitude. (**a**) Illustration of experimental apparatus and task structure for Exp. 1. (**b**) Schematic view of clamped visual feedback paradigm, in which the angular path of the cursor is independent of hand movement direction. (**c-d**) Behavior for all groups (n = 12/group), divided into two panels for visualization purposes. The small clamp groups (**c**) demonstrate adaptation rates which scale with error size, whereas the large clamp groups (**d**) show saturated responses. (**e**) Segmented regression indicates that the initial adaptation rate scales between 0° and 4.4° before saturating for all errors above this breakpoint (dashed vertical lines represent 95% C.I.). (**f**) Sensorimotor aftereffects, measured during the first cycle following the clamp block. Dots are individuals; shading and error bars denote SEM. Gray shading denotes cycles without visual feedback.

The use of clamped visual feedback offers a new tool to address a fundamental problem in error-based learning, namely, how does the response of the system vary as a function of error size? In studies using a fixed, task-relevant perturbation (e.g., standard visuomotor rotation), error size is confounded with learning: As learning unfolds, the mean error size becomes dramatically smaller. To provide a cleaner assay of the responsiveness of the adaptation system to errors of varying size, previous studies have used errors that vary randomly in size and direction from trial to trial, such that the overall mean error is zero^10,14,15^. A limitation with this approach is that one can only measure trial-by-trial changes. In contrast, with the clamp method, we can examine the full accumulated adaptive response to errors of a given size since the error signal remains invariant. Thus, we can assess not only how changes in error magnitude influence the response of the system, but also how this responsiveness might change over time and training.

Our initial study with clamped visual feedback revealed learning functions that were surprisingly invariant over a wide range of error sizes (7.5° – 95°)^17^. This invariance was evident in the initial rate of adaptation as well as in the final asymptotic value. As noted above, prior studies indicate that sensitivity is reduced to large errors^8,10,14,15^; it may be that the smallest value previously tested with the clamp method (i.e., 7.5°) falls within the range in which the error-driven response is already saturated.

In the current study, we focus on small clamped errors (i.e., errors < 7.5°), using perturbation sizes that are more representative of the feedback that we typically experience from intrinsic motor variability^18^. We expect that the response to these smaller clamped errors will be dependent on the size of the error, and thus, allow us to estimate the saturation point. Assuming we observe some scaling of the response as a function of error size, the clamp method also allows us to ask if this is evident in both the learning rate and asymptote as predicted by current models of adaptation.

## Results

### Initial adaptation rates only scale with error size for small errors

In a between-subject design, participants (n = 96, 12/group) were presented with visual feedback that was clamped to a fixed path which was angularly offset from the target by 0°, 1°, 1.75°, 3.5°, 6°, 10°, 15°, or 45°. This manipulation was explicitly described to the participants and they were instructed to ignore the feedback and simply move directly to the target (Fig. 1a). With the exception of the 0° control group, all groups implicitly adapted to the clamp (t_11_ =-.04, p =. 97 for 0° group; t_11_>5.9, p<.0001 for all other groups; Fig. 1c,d).

Within all adapting groups, there was an effect of clamp size on the average per-trial rate of learning over the first five cycles (ANOVA: F_6,77_ = 6.45, p<.0001, η^2^ =. 33). Although there was a modest linear relationship between clamp size and early adaptation rate (r_82_ =. 29, p = 0.01), the adaptation rate appeared to be composed of two zones, one where the rate scaled in proportion to error size, and another where rates were invariant. To formally assess this hypothesis, we performed segmented linear regressions. Taking model complexity into account, a two-region segmented regression yielded the best model (see Supplementary Fig. 1). This model predicted that the breakpoint between the proportional and saturated zones was at the remarkably low value of 4.4° (95% CI [3.9°, 4.9°], Fig. 1e).

These results, in combination with previous work^8,10,14,15,17^, are clearly at odds with models entailing a fixed learning rate (i.e., adaptation scaling linearly with error size). Prior observations of a nonlinear response to error have inspired models in which the learning rate saturates for large errors^13^, or large errors are discounted before the update step of the learning process^10,14,19^. If exposed to a constant perturbation, these models can generate similar adaptation rates and asymptotes in response to large errors. However, these models would also predict a lower asymptote in response to small errors in the linearly proportional zone. Contrary to this prediction, with the exception of the 1° group, the magnitude of adaptation at the end of training was similar for a wide range of clamp offsets (Fig. 1f and Supplementary Note 1).

### Adaptation converges on a common asymptote

The observation of similar performance across a range of error sizes at the end of training is tempered by the fact that we did not have a sufficient number of trials to ensure that learning had become asymptotic; as such, it is unclear if prolonged exposure to constant errors of varying size will converge at a common asymptote. To address this issue, we conducted a second experiment in which the number of cycles was increased from 40 to 160. Participants (n = 30, 10/group) were exposed to clamped feedback with an angular offset of 1.75°, 3.5°, or 15°. These offsets were chosen because they span the range of early adaptation rates observed in Experiment 1. Consistent with the results of Experiment 1, there was a clear scaling of the rates across the proportional zone (ANOVA: F_2,27_ = 18.6; p<.0001; η^2^ =. 58), with Tukey Kramer post hoc tests revealing significant differences between all pairwise comparisons (Fig. 2a). Strikingly, the three groups reached a similar asymptote, with all groups demonstrating final aftereffects of ∼25° (ANOVA: F_2,27_ = 0.39, p =. 68; η^2^ = 0.03) (Fig. 2b; see also Supplementary Note 2).

**Figure 2.**
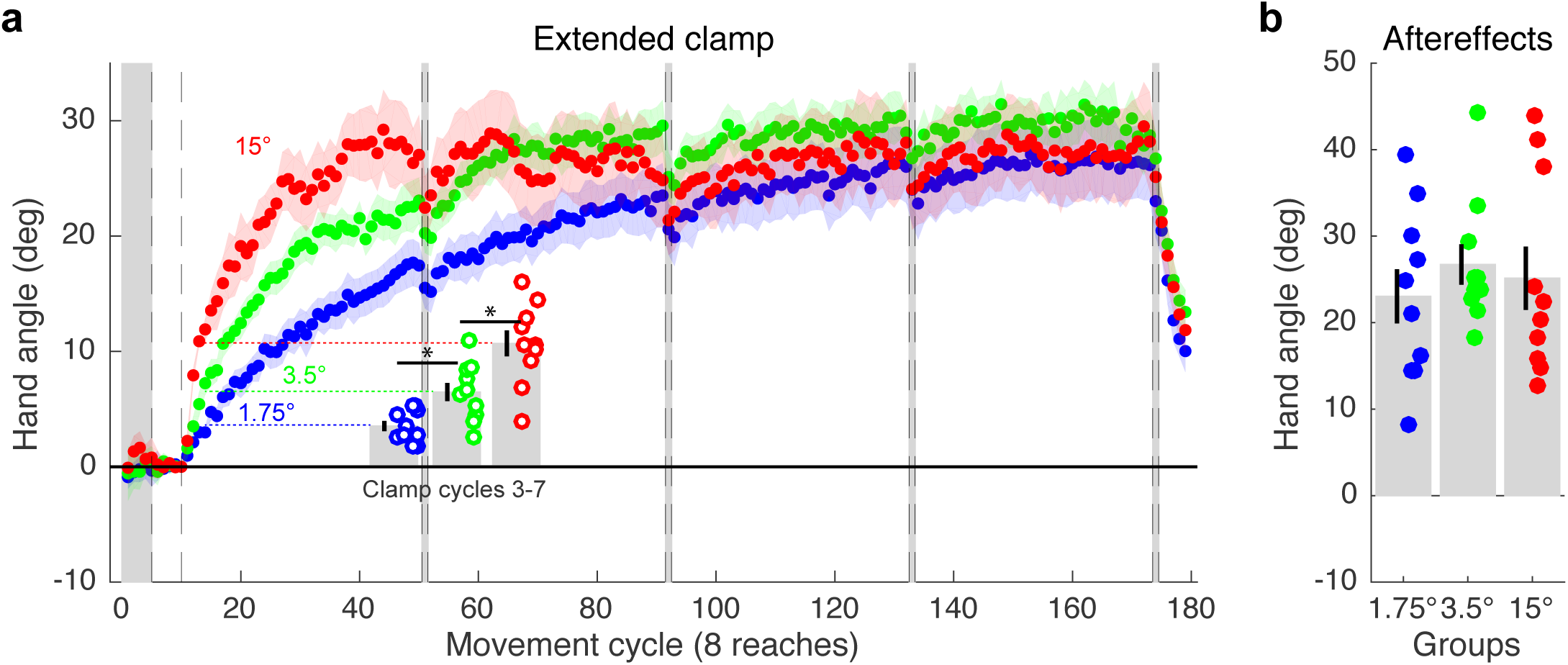
Implicit adaptation converges on a common asymptote. (**a**) The 1.75°, 3.5°, and 15° clamp groups in Exp. 2 (n = 10 / group) adapted at markedly different rates (bar graphs depict mean of cycles 3-7). However, there was convergence of all three learning functions by the end of 160 cycles, and (**b**) no difference between groups in the size of the final aftereffects. Asterisk in (**a**) denotes significant differences between groups early in the clamp phase. Dots are individuals; shading and error bars denote SEM.

## Discussion

This dissociation between size-dependent early adaptation rates and invariant asymptotic adaptation is at odds with models that correct for a constant fraction of error size as well as error discounting models (See Supplementary Fig. 2 and Supplementary Note 3). Even if the learning rate varies as a function of error size, assuming a fixed retention factor, these models predict that asymptotic behavior will diverge since the asymptote is determined by the equilibrium between learning and forgetting.

In addition to identifying a fundamental limitation with current models of sensorimotor adaptation, our results draw attention to a more general issue. Behavioral responses to error are usually interpreted through the lens of error sensitivity. This perspective is apparent not only in studies of visuomotor adaptation, but is also evident in research on saccadic^20^, locomotor^21^, and force field adaptation^22^. The sensitivity function is generated by dividing the magnitude of the motor correction by the error size. When applied to the behavioral data that we and others have observed, this divisive operation generates a function in which sensitivity is high for small errors and gradually decreases to near zero for large errors (Fig. 3a,c).

**Figure 3.**
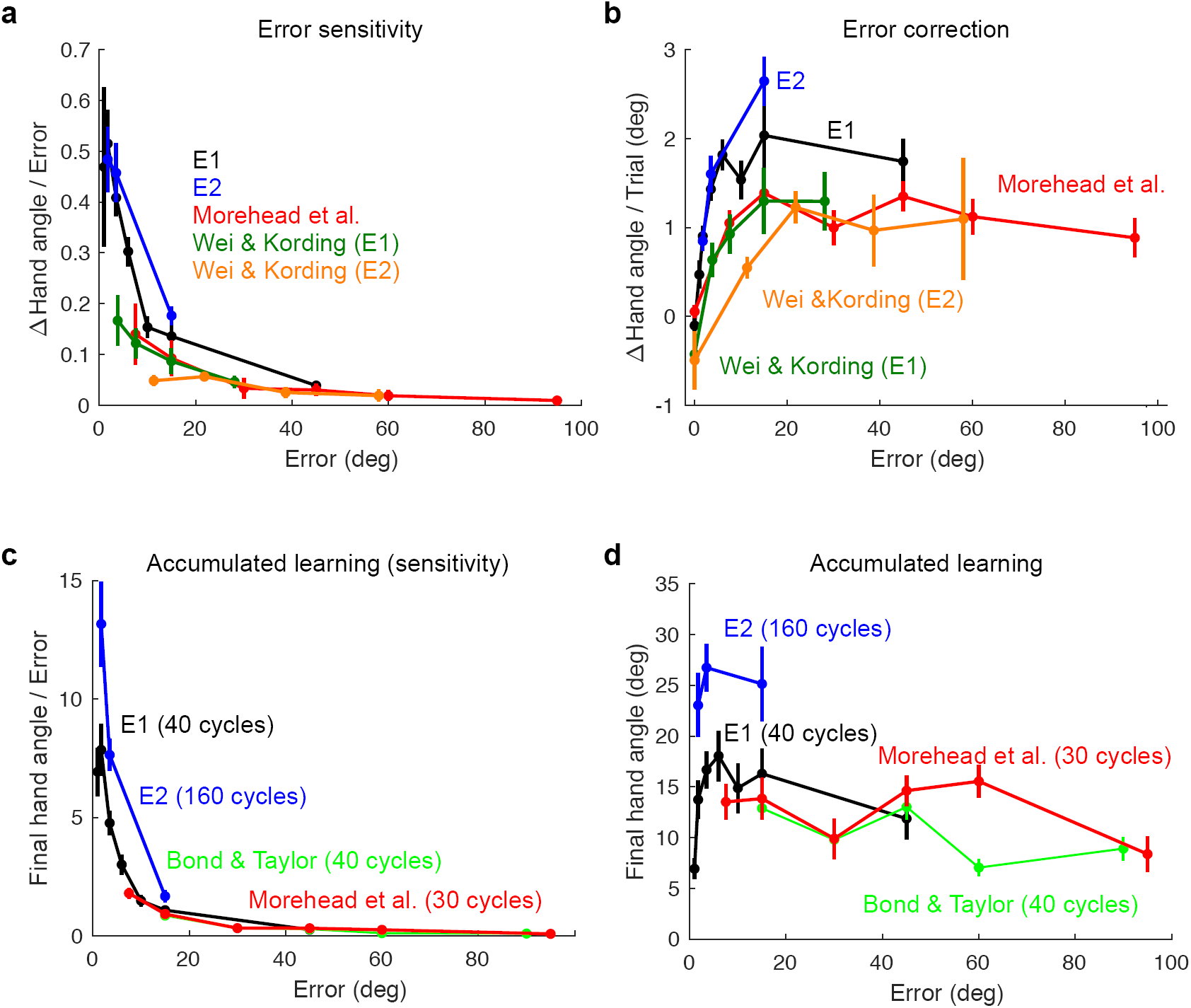
Adaptation assessed in terms of error sensitivity (left) or error correction (right). Here we plot data from several different studies^10,17,29^, including the present one, using two ways to consider trial-by-trial changes in hand angle as a function of error size (see Methods). (**a**) Error sensitivity, operationalized as the change in hand angle divided by error size, starts at an early maximum and quickly decays as errors increase in size. **(b)** The same data, plotted in terms of the untransformed error correction, shows a function that starts small and then saturates, suggesting that the motor system continues to produce a robust, invariant response over a wide range of error sizes. Plotting the aftereffect data in terms of a sensitivity function (**c**) also fails to capture the relative invariance of these data within a given experimental context (**d**). Note the one discrepant point from Exp. 1 in panels C and D from the 1° clamp condition; we suspect this is due to an insufficient number of trials to approximate asymptotic performance. Error bars denote SEM.

Although this error sensitivity metric is mathematically capable of approximating the behavioral effects observed with variation in error size, focusing on the untransformed behavioral responses to errors of varying size suggests a different perspective on the limiting factor in adaptation. As seen in Fig. 3b, adjustments in motor output scale for small errors before quickly reaching a saturation point that holds across a broad range of larger errors (Fig. 3b). Depicting the actual behavioral change from sensory prediction error highlights the limited dependency of the system on error size, as well as the common asymptotic level of learning in response to small and large errors (Fig. 3d). Thus, incorporating an update rule in which the correction (i.e., behavioral change), rather than error sensitivity, is modeled as a function of error size may offer a more appropriate framework for understanding the constraints underlying sensorimotor adaptation. The data in Fig. 3b suggest that the error correction function, expressed in terms of absolute change in heading direction from trial-to-trial, would have a half-sigmoid shape with a saturation point at a small error size.

We can envision three, non-mutually exclusive ways in which current models of adaptation could be modified to capture size-dependent early adaptation rates for small errors combined with invariant asymptotic adaptation. First, the learning rate and retention parameters could be coupled, scaling together with error size^23^. For instance, small errors may elicit smaller corrections and greater retention, while large errors may elicit larger corrections but weaker retention. Whereas current models have considered that learning rate may be dependent on error size, this variant would require that the forgetting process is also error-size dependent (the *A* term in the state-space model equation). Moreover, to achieve a common asymptote across different error sizes constrains the form of the coupling between these two parameters.

Second, the adaptation system may normalize its responses to sensory prediction errors with repeated exposure, akin to normalization processes observed in response to reward prediction errors^24^. For example, the system may increase its responses to small, yet persistent errors. Alternatively, responses to large errors may diminish over time until reaching some intermediate normalized update size. By this normalization hypothesis, the size of the motor correction changes over trials and, due to the invariance of the size of the clamped error, eventually converges on the same value for all error sizes.

To this point, we have assumed that the clamped visual error is the primary signal driving the change in behavior; however, other error signals, in particular signals arising from proprioception, may also impact adaptation to a visual perturbation. Thus, a third possibility is that the asymptotic response may reflect the limit of proprioceptive recalibration, which is independent of visual error size. That is, as the heading angle changes due to the clamped visual error, the proprioceptive sensory prediction error would increase, but with the opposite sign. The asymptote would correspond to the balance point between these two opposing error signals.

Future work will be required to formalize these hypotheses and develop experimental tests to evaluate the different mechanisms. Regardless of the appropriate reformulation of models of sensorimotor adaptation, we expect it will be fruitful to shift the focus away from the error sensitivity of the learning system, and instead, address the constraints on the behavioral change that arises in response to the error.

## METHODS

### Participants

Healthy, young adults (N=126, 89 females, age = 21±2 years old) were recruited from the University of California, Berkeley, community. Each participant was tested in only one experiment. All participants were right-handed, as verified with the Edinburgh Handedness Inventory^25^. Participants received course credit or financial compensation for their participation. The Institutional Review Board at UC Berkeley approved all experimental procedures.

### Experimental Apparatus

The participant was seated at a custom-made tabletop housing an LCD screen (53.2 cm by 30 cm, ASUS), mounted 27 cm above a digitizing tablet (49.3 cm by 32.7 cm, Intuos 4XL; Wacom, Vancouver, WA). The participant made reaching movements by sliding a modified air hockey “paddle” containing an embedded stylus. The position of the stylus was recorded by the tablet at 200 Hz. The experimental software was custom written in Matlab, using the Psychtoolbox extensions^26^.

### Reaching Task

Center-out planar reaching movements were performed from the center of the workspace to targets positioned at a radial distance of 8 cm. Direct vision of the hand was occluded by the monitor, and the lights were extinguished in the room to minimize peripheral vision of the arm. The start location and target location were indicated by white and blue circles, respectively (both 6 mm in diameter).

To initiate each trial, the participant moved the digitizing stylus into the start location. The position of the stylus was indicated by a white feedback cursor (3.5 mm diameter). Once the start location was maintained for 500 ms, the target appeared at one of 8 locations, placed in 45° increments around a virtual circle. Participants were instructed to accurately and rapidly “slice” through the target, without needing to stop at the target location. Visual feedback, when presented, was provided during the reach until the movement amplitude exceeded 8 cm. As described below, the feedback either matched the position of the stylus (veridical) or followed a fixed path (clamped). If the movement was not completed within 300 ms, the words “too slow” were generated by the sound system of the computer.

In Experiment 1, the position of the cursor was frozen for 1 s once the movement amplitude reached 8 cm. The participant was free to begin moving back to the start location during this time. After the spatial feedback period, the cursor disappeared. Once the participant’s hand was back within 2 cm of the start circle, a white ring appeared, indicating the radial distance between the hand and center start position. The ring was displayed to aid the participant in returning to the start location, without providing angular information about hand position. Two changes were made in Experiment 2: First, the cursor was turned off 50 ms after the hand crossed the virtual target ring. Second, during the return movement, the feedback cursor reappeared when the participant’s hand was within 1 cm of the start. These changes reduced the time required for each trial and allowed the participants to complete the extended number of trials required in Experiment 2 within our time constraints. Average total trial time in Experiment 1 was 4.45 ± .62 s versus 2.51 ± .26 s in Experiment 2.

### Experimental Feedback Conditions

Across the experimental session, there were three types of visual feedback. On no-feedback trials, the cursor disappeared when the participant’s hand left the start circle and only reappeared at the end of the return movement. On veridical feedback trials, the cursor matched the position of the stylus during the 8 cm outbound segment of the reach. On clamped feedback trials, the feedback followed a path that was fixed along a specific heading angle^17,27,28^. The radial distance of the cursor from the start location was still based on the radial extent of the participant’s hand during the 8 cm outbound segment, but the angular position was fixed relative to the target (i.e., independent of the angular position of the hand).

The primary instructions to the participant remained the same across the experimental session: Specifically, that they were to reach directly towards the visual target. Prior to the introduction of task-irrelevant clamped feedback trials, participants were briefed about the feedback manipulation. They were informed that the position of the cursor would now follow a fixed trajectory and that the angular position would be independent of their movement. They were explicitly instructed to ignore the cursor and continue to reach directly to the target. The same instructions in abbreviated form (“Ignore the cursor and move your hand directly to the target location”) were repeated verbally and with onscreen text after 20 movement cycles in Experiment 1 (exact mid-point) and every 40 movement cycles during Experiment 2.

#### Experiment 1

In a previous experiment, adaptation to task-irrelevant clamped visual feedback was statistically uniform to offsets between 7.5°-95°. The main goal of Experiment 1 was to investigate if there was a dependency on error size for angles smaller than 7.5°. Participants (n = 96, 12/group) were randomly assigned to one of eight groups that differed in terms of the size of the clamped visual feedback: 1°, 1.75°, 3.5°, 6°, 10°, 15°, and 45° (with a 0° group included as a control). The Euclidean distances for these clamp offsets, measured from the centers of cursor and target, were as follows (smallest to largest, in mm): 0, 1.4, 2.4, 4.9, 8.4, 13.9, 20.9, and 61.2. Given that the target diameter was 6 mm and the feedback cursor diameter was 3.5 mm, a substantial portion of the cursor overlapped with the target for the 1° and 1.75° clamps, and was fully embedded in the case of the 0° clamp. Half of the participants trained with a clockwise clamp offset, and the other half with a counterclockwise clamp offset.

The session began with two baseline blocks, the first comprised of 5 movement cycles (40 reaches to 8 targets) without visual feedback and the second comprised of 5 cycles with a veridical cursor displaying hand position. The experimenter then informed the participant that the visual feedback would no longer be veridical and would now be clamped at a fixed angle from the target location. The clamp block had 40 cycles. A short break (<30 s), as well as a reminder of the task instructions, was provided at the mid-way point of this block. Immediately following the perturbation block, there were two washout blocks, first a 5 cycle block in which there was no visual feedback, followed by 5 cycles with veridical visual feedback. Participants were debriefed at the end of the experiment and asked whether they ever intentionally tried to reach to locations other than the target. All subjects reported aiming to the target throughout the experiment.

#### Experiment 2

In Experiment 2 we assessed adaptation over an extended number of task-irrelevant clamped visual feedback trials. The purpose of extending the perturbation block was to ensure that participants reached asymptotic levels of learning. We were particularly interested in whether asymptotic adaptation would converge in response to small and large clamps.

Participants (n = 30, 10/group) were assigned to either a 1.75°, 3.5°, or a 15° clamped visual feedback group. Clockwise and counterclockwise perturbations were counterbalanced within each group. As in Experiment 1, the session started with two baseline blocks, 5 cycles without visual feedback and then 5 cycles with veridical feedback. However, the number of trials in the clamped visual feedback block was quadrupled to 160 cycles. We included 1 cycle with no visual feedback after every 40 movement cycles. The purpose of these interspersed no-feedback trials was to gauge adaptation magnitudes in the absence of the learning stimulus (i.e., clamped visual feedback) at different time points within the extended clamp block (see Supplementary Figure 3). Immediately prior to the no feedback block, the participant was informed that there would be a few trials without feedback and reminded to always reach directly to the target. The experiment ended with a final block of 5 cycles with veridical visual feedback of the participant’s hand position.

### Comparison of error sensitivity and error correction

For the comparison of error sensitivity and error correction functions in Figure 3, we used the early adaptation rate (panels a and b) and aftereffect data (panels c and d) from the present study, as well as data sets from three other studies that have compared adaptive responses to a range of error sizes^10,17,29^. The data from Experiments 1 and 2 in Wei and Kording were transformed from Cartesian coordinates (as presented in their paper) to polar coordinates. The data from Bond and Taylor were restricted to the aftereffect data from their Experiment 3 (comparison of adaptation to different rotation sizes). We used the data from the initial adaptation cycles (mean change in hand angle over first ten movement cycles) and aftereffect phase from Experiment 4 of Morehead et al. (comparison of different clamp offsets). To obtain measures of error sensitivity, the raw response magnitudes were divided by their corresponding error size. We did not include the data from the 1° group in our Experiment 1 since the group function was clearly not at (or likely approaching) asymptote.

### Data Analysis

All statistical analyses and modeling were performed using Matlab 2015b and the Statistics Toolbox. The primary dependent variable in all experiments was endpoint hand angle, defined by the angle of the hand position relative to the target at the time the radial distance of the hand reached 8 cm from the start position (i.e., angle between lines connecting start position to target and start position to hand). Additional analyses were performed using hand angle at peak radial velocity rather than endpoint hand angle. The results were essentially identical for the two dependent variables; as such, we only report the results of the analyses using endpoint hand angle.

Outlier responses were removed from the analyses. To identify these, the Matlab “smooth” function was used to calculate a moving average (using a 5-trial window) of the hand angle data for each target location. Outliers were trials in which the observed hand angle deviated by more than 3 standard deviations from the moving average function. This procedure resulted in the elimination of ∼1% of trials involved in our statistical analyses of early adaptation rates and aftereffects; our findings are the same whether tests were performed with or without outlier removal (values reported in main text are with outlier removal). In total, less than 1% of trials overall, with a maximum of 2% for an individual, were removed.

Movement cycles consisted of 8 consecutive reaches (1 reach/target). Early adaptation rate was quantified by averaging the endpoint hand angle values over cycles 3-7 of the clamp, and dividing by the number of cycles (i.e., 5) to get an estimate of the per trial rate of change in hand angle. (As a check, we performed a secondary analysis using cycles 2-10 and obtained nearly identical results.) We opted to use this measure of early adaptation rather than obtain parameter estimates from exponential fits since the latter approach gives considerable weight to the asymptotic phase of performance and, therefore would be less sensitive to early differences in rate. This would be especially problematic in Experiment 2. The aftereffect was quantified by using the data from the first no-feedback cycle following the last clamp cycle. Details for all four no-feedback cycles in Experiment 2 are provided in the Supplemental section.

All t-tests were two-tailed. In order to confirm that there was a robust adaptive response in Experiment 1, a paired t-test was performed comparing baseline hand angle during the last cycle of the veridical feedback baseline to the first no feedback cycle (i.e., aftereffect) immediately following the perturbation block. Posthoc tests following significant ANOVAs were performed using Tukey-Kramer’s Honest Significant Difference in order to determine specific differences in group means. Partial eta squared (?^2^) values are provided as a measure of effect size.

For the segmented linear regression (SLR) performed in Experiment 1, contiguous regression lines were fit to the data, with each line having an independent intercept and slope. Parameters for the regression lines were identified by a least-squares fitting procedure. Boundaries for the adjoining segments were estimated by finding the break point(s), an additional parameter defining where two separate regression lines meet, that minimized the residual sum of squares. Relative fits were compared using corrected Akaike Information Criterion (AICc) values, a procedure that adjusts for the number of data points and assigns penalties for extra parameters.

No statistical methods were used to predetermine sample sizes. The chosen sample sizes were based on our previous study using the clamp method^17^, as well as prior psychophysical studies of human sensorimotor learning^28,30–32^.

### Reaction and Movement Times

Average movement times were quite fast, averaging 119 ± 29 ms in Experiment 1 and 136 ± 25 ms in Experiment 2. The instructions did not impose any constraints on reaction time. On average, in Experiment 1 participants initiated their reaches in 429 ± 72 ms, while in Experiment 2 reaction times were 364 ± 54 ms. No significant correlations were found between these temporal variables and our primary measures of adaptation (rate and aftereffect magnitude).

#### Code and Data Availability

All code and data are available upon request.

#### Author contributions

*HEK*: Conceptualization; Data curation; Software; Formal analysis; Investigation; Visualization; Methodology; Writing — original draft; Writing — review and editing

*JRM*: Conceptualization; Formal analysis; Methodology; Writing — review and editing

*DEP*: Conceptualization; Formal analysis; Writing — review and editing

*RM*: Formal analysis; Methodology; Writing — review and editing

*RBI*: Conceptualization; Formal analysis; Supervision; Funding acquisition; Writing — original draft; Writing — review and editing

## Acknowledgments

We thank Wendy Shwe for assistance with the data collection. We are also grateful to Maurice Smith and Matthew Boggess for helpful discussions. This work was supported by grant NS092079 from the National Institutes of Health.

## Competing Interests

No competing interests, financial or otherwise, are declared by the authors.

## SUPPLEMENTAL INFORMATION

### Supplementary Note 1

#### Aftereffect data

In Experiment 1, there was an effect of error size on the aftereffect data obtained after 40 cycles of exposure to the clamped feedback (F_6,77_ = 3.25, p =. 007, η^2^ =. 20, Fig. 1g). However, this effect was driven by the 1° group, and likely due to the trivial reason that the design did not include a sufficient number of training cycles for this group to reach asymptote. A Tukey-Kramer posthoc test revealed the only significant differences were between the 1° group and larger clamp sizes. Furthermore, excluding the 1° group, there were no reliable differences among all of the other pairwise comparisons (all t_22_<2.07, all p>.05). Indeed, the magnitude of the aftereffect was not correlated with error size (r_82_ = 0.04, p = 0.71). These results suggested that asymptotic adaptation may be independent of error size, a hypothesis which was more rigorously tested in Experiment 2.

### Supplementary Note 2

#### Comparisons between Experiment 1 and Experiment 2

Overall, the magnitudes of final aftereffects in Experiment 2 were considerably larger than the aftereffect of matched groups in Experiment 1. One reason for this is, of course, the increase in the number of clamp cycles in Experiment 2, which had 160 compared to the 40 cycles used in Experiment 1. As can be seen in Supplementary Figure 3, the magnitude of adaptation is still rising after 40 cycles of clamped feedback. As an exploratory analysis, we performed a between-experiment 2-way ANOVA, with one factor being experiment (1 or 2) and the other error size, limiting this to the three conditions in Experiment 1 that were included in Experiment 2 (1.75°, 3.5° and 15°). We compared the aftereffect data after 40 clamp cycles (i.e., final aftereffects from Experiment 1 compared to first no feedback probe in Experiment 2). There was a significant effect of experiment (F_1,60_ = 5.27, p =. 03, η^2^ =. 07), a marginal effect of clamp size (F_2,60_ = 2.8, p =. 07, η^2^ =. 08), and no interaction (F_2,60_ =. 57, p =. 57, η^2^ =. 02). These results suggest that some performance differences may have arisen from the methodological changes introduced in Experiment 2, namely, eliminating frozen endpoint feedback and utilizing a quicker method for finding the start position before each trial (see Methods). This resulted in a shorter total trial duration in Experiment 2, likely reducing the decay of learning associated with delay^16^. This point highlights that asymptotic magnitude is context specific, as adaptation is highly sensitive to variables such as temporal delay between hand and cursor movement^17,18^, viewing angle^19^, and the nature of visual feedback (e.g., online versus endpoint feedback)^20,21^. As such, our claim that the asymptote is independent of error size holds for a given context; the value will shift in other contexts, although the shift will be uniform for all error sizes.

### Supplementary Note 3

#### Computational models of sensorimotor adaptation predict divergence of asymptotic adaptation when early adaptation rates are different

A single rate state-space model can generally provide a reasonable account of the performance changes observed in standard adaptation studies in which the feedback is contingent on the movement^4–6^. This model takes the following form:

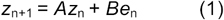

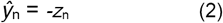

*z*_*n*_ represents the state estimate of the perturbation on trial *n*. The learning rate, *B*, corresponds to the proportion of the error, *e*_*n*_, that is corrected for, and *A* is a retention factor which represents the proportion of the state retained from one trial to the next. The reach direction relative to the target on trial *n, ŷ*_*n*,_ is opposite in sign to the state estimate. Note that unlike standard adaptation experiments in which the error changes over trials, the error remains constant with the clamp method.

We evaluated different state-space models, focusing on the data from Experiment 2 (reproduced in Supplementary Fig. 2a). For our model fitting and simulation procedures we applied standard bootstrapping techniques, constructing group-averaged hand angle data 1000 times by randomly resampling with replacement from the participant pool. Using Matlab’s *fmincon* function, we estimated the retention and learning parameters which minimized the least squared error between the bootstrapped data and model output (*ŷ*_*n*_). All values in brackets represent the 95% bootstrapped C.I. of the resampled means.

We first verified that a single rate state-space model, which in its basic form assumes single, constant *A* and *B* values for all error sizes, cannot account for the key features of the current experiments. We estimated *A* and *B* for each of the 1000 bootstrap simulations ([.86,. 95] and [.11,. 29] for *A* and *B*, respectively) (Supplementary Fig. 2b). In contrast to the invariant asymptotes observed in Experiment 2, the simulated functions generated with these parameter estimates showed wildly different asymptotic values: [3.2°, 4.8°], [6.5°, 9.6°], and [27.9°, 41.0°] in response to 1.75°, 3.5°, and 15° clamps, respectively. As the error term is constant during a clamp (i.e., e_n_ is equal to clamp size on every trial), and the asymptote is equal to *(Be)/(1-A)*, different error terms (i.e., clamp sizes) will always generate different asymptotic values if *A* and *B* are both fixed.

An alternative class of models posit that performance changes reflect the activity of multiple learning processes, each operating in a similar manner but over different time scales^7,8^. For example, in a dual-rate state-space model of adaptation^8^, the output (5) is the sum of two single rate models, one in which both learning and forgetting occur over a faster time scale (3) than the other (4). Despite this added flexibility, this model is similarly constrained as the single rate model. As previously explained, the asymptotes for each state will still be equal to *(Be)/(1-A)*, meaning different size clamps eventually reach different asymptotes. Indeed, our simulations show that a dual rate state-space model will generate functions (Supplementary Fig. 2c) which are qualitatively similar to the single rate model (Supplementary Fig. 2b).

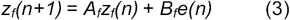

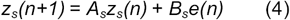

*where A*_*f*_ *< A*_*s*_, *B*_*f*_ *> B*_*s*_

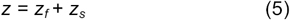

As noted in the main text, various studies have suggested that the learning rate, *B*, may vary with error size^9–14^. To examine this class of models, we used a single rate state-space model (eqns. 1 and 2) with a common retention factor and three learning rates, one for each error (clamp) size. We again obtained 1000 sets of parameter estimates from the bootstrapped data in order to generate the simulations seen in Supplementary Fig. 2d. This 4-parameter model (a common *A*, and separate *B* values for each clamp size) also failed to capture the key features of the behavior. Although the model generated more similar asymptotes across clamp sizes than the single rate state-space model with constant *A* and *B* ([18.4°, 28.8°], [25.1°, 32.6°] and [21.6°, 34.5°] for 1.75°, 3.5°, and 15° clamps), it failed to capture the clear separation of early adaptation rates observed in the behavioral data (Supplementary Fig. 2a). In fact, the model predicted nearly identical early adaptation rates (i.e., mean change in hand angle per movement cycle over first 5 cycles) for the 3.5° and 15° clamps ([.9°, 1.4°] and [.7°, 1.5°]), and only a modestly slower rate for the 1.75° clamp [.7°, 1.1°]. Moreover, *B* values in the adaptation literature^4,5,15^ typically range from. 10 to. 30, whereas the *B* values required to fit the data for the small error clamps would have to go as high as. 70, and even then, provide a relatively poor match to the observed behavior.

We also considered an alternative means of parameter estimation with this model (eqns. 1 and 2) in which we recapitulate the different adaptation rates by using the best fit *A* value of. 95 and approximating the mean *B* values directly from the early cycles of the bootstrapped data, using the mean change in hand angle over the first 5 cycles divided by clamp size. This procedure yielded the following *B* values: [.36,. 60], [.31,. 59], and [.13,. 21] for 1.75°, 3.5°, and 15° clamps, respectively. Here the predicted asymptotes in response to the three clamp sizes again strayed far from the observed behavior: [12.2°, 20.1°], [20.4°, 39.3°], and [36.4°, 60.4°].

Lastly, we tested a model^13^ motivated by observations suggesting that the perceived error is linear for small errors before saturating at an upper bound for larger errors^9^. To model this nonlinear effect, the authors replaced the error term, *e*, with a function which saturates for larger errors (*s*tanh(e*_*n*_*/s*)). *s* is a scaling parameter for the hyperbolic tangent function:

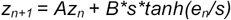

This model produced learning functions which suffered from the same shortcomings as the state-space model with error-dependent learning rates, again predicting virtually identical early adaptation rates for the 3.5° and 15° clamp ([.8°, 1.3°] and [.8°, 1.5°]), a *B* value [.70, 1.0] which far exceeds normal estimates of this value, and a slightly lower asymptote for the 1.75° clamp ([18.7°, 27.4°] versus [24.8°, 31.6°] and [25.2°, 33.7°] for the 3.5° and 15° clamps, respectively).

In summary, even models that account for variation in sensitivity to error fail to capture the combined effects of divergent early adaptation rates and invariant asymptotic performance (see Moazzezi 2018 for a comprehensive analysis of this issue)^22^.

## Figure Legends

**Figure S1.**
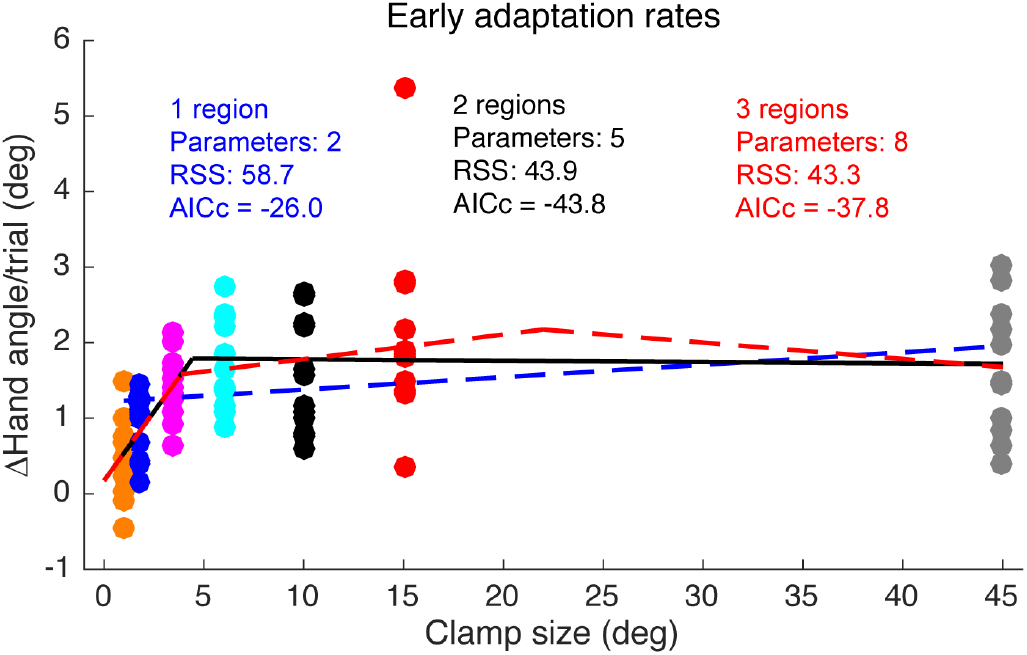
Segmented regression analysis of early adaptation rates. In our previous study using the clamped feedback method, we observed similar adaptation rates for clamp offsets of 7.5°-95°. In the current study, we focus on clamps < 7.5°. As described in the text, there was a clear scaling of adaptation rates for the smallest clamps (Figs. 1e, 2a). To determine the error size at which adaptation rates saturate (i.e., the breakpoint), we performed a segmented regression analysis, a widely used method for fitting data that are hypothesized to have at least two different linear regions^3^. This figure shows the results for a simple linear regression (1 region) and two segmented regression models, one with 2 regions and the other with 3 regions, along with the residual sum of squares (RSS) and corrected Akaike Information Criterion (AICc) scores. The best model, as determined by the AICc scores, was the two-region model. Models with more than three regions scored progressively worse and are not shown. Consistent with our prediction of a breakpoint < 7.5°, the predicted breakpoint was 4.4°.

**Figure S2.**
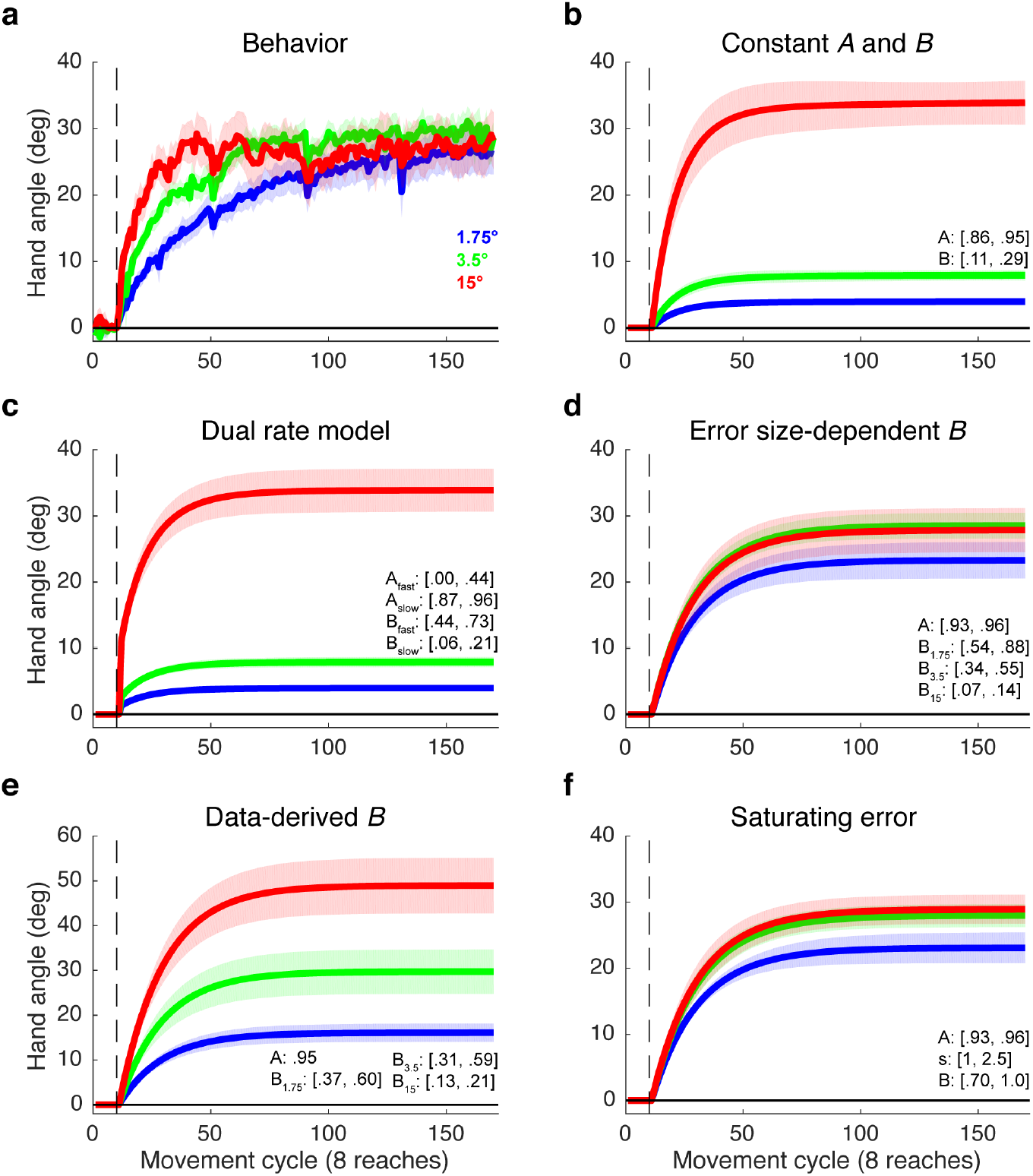
Existing models fail to capture both early learning and asymptotic behavior observed during adaptation to different error sizes. (**a**) Behavioral data from Experiment 2 (same as Fig. 2 of main text, but with no-feedback cycles removed). (**b**) Simulation of a single rate state-space model with a common retention factor and learning rate (best fit *A* and *B* of bootstrapped group means). **(c)** Simulation of a dual-rate model, with separate retention and learning rate parameters that operate over different time scales (e.g., fast and slow). (**d**) Simulation using a common retention factor and error size-dependent learning rates (best fit *A* and learning rates, *B*_*e*_, for each clamp size). The simulated functions predict indistinguishable adaptation rates for the 3.5° and 15° clamp conditions over the initial portion of the perturbation block, in contrast to the markedly different rates observed in the actual behavior. **(e)** Simulation using estimates of *B* derived from the behavioral data. The functions approximate the early separation between learning functions, but predict divergent asymptotes. (Note: y-axis scaling was changed to fit data). **(f)** Based on a model from Tanaka et al. (2012), a saturating error function can capture the nonlinear effects of error size on adaptation. However, similar to the fits in ***d***, this model also predicts similar early adaptation rates for the three error sizes, as well as a lower asymptote for the 1.75° clamp. 95% CIs for parameter estimates in brackets. Lines and shading denote mean and SEM, respectively.

**Figure S3.**
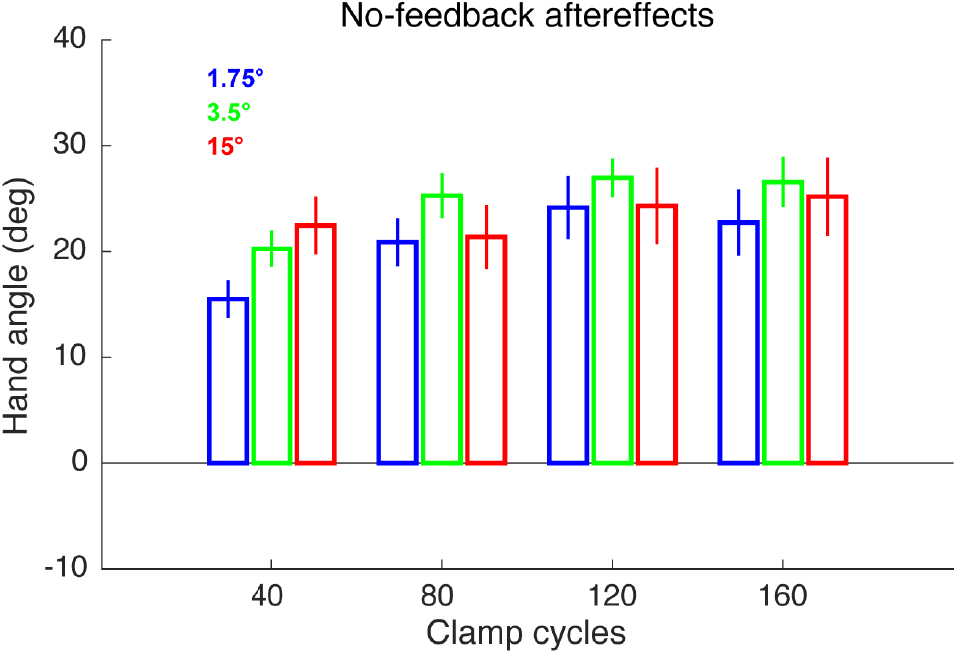
Data for all of the no-feedback probes (x-axis represents number of clamp cycles preceding the no-feedback probe) from Experiment 2. The results of a two-way repeated measures ANOVA revealed no main effect for group (between subjects factor: F_2,27_ =. 62, p =. 54, η^2^ =. 04), a significant effect for probe (within subjects factor: F_3.81_ = 19.2, p<10^−7^, η^2^ =. 07), and a significant interaction effect (F_6,81_ = 2.64, p =. 02, η^2^ =. 02). Due to the interaction, separate between subjects one-way ANOVAs were conducted for each of the four no feedback cycles. There was a marginal effect of group (i.e., clamp size) at the first probe only (F_2,27_ = 2.77; p =. 08; η^2^ =. 21). Over subsequent blocks learning functions for each group began to converge, resulting in the corresponding values for no-feedback probes 2-4: F_2,27_ =. 92,. 30,. 39; p =. 41,. 75,. 68; η^2^ =. 07,. 02,. 03).

